## Supplemental Figurres for "Virus specificity and nucleoporin requirements for MX2 activity are affected by GTPase function and capsid-CypA interactions"

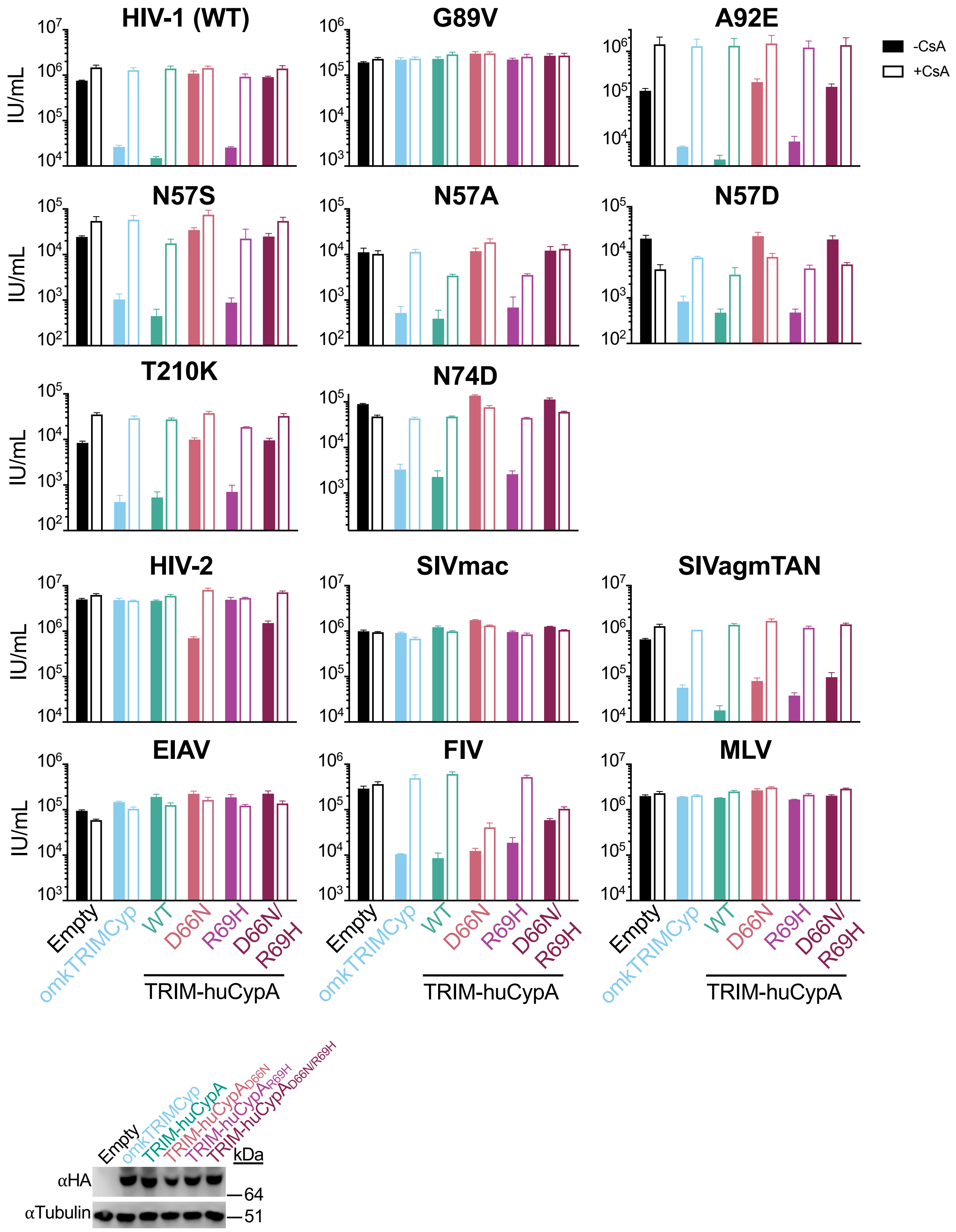


**S1 Fig. Recognition of retroviral capsids by TRIMCyp fusions**

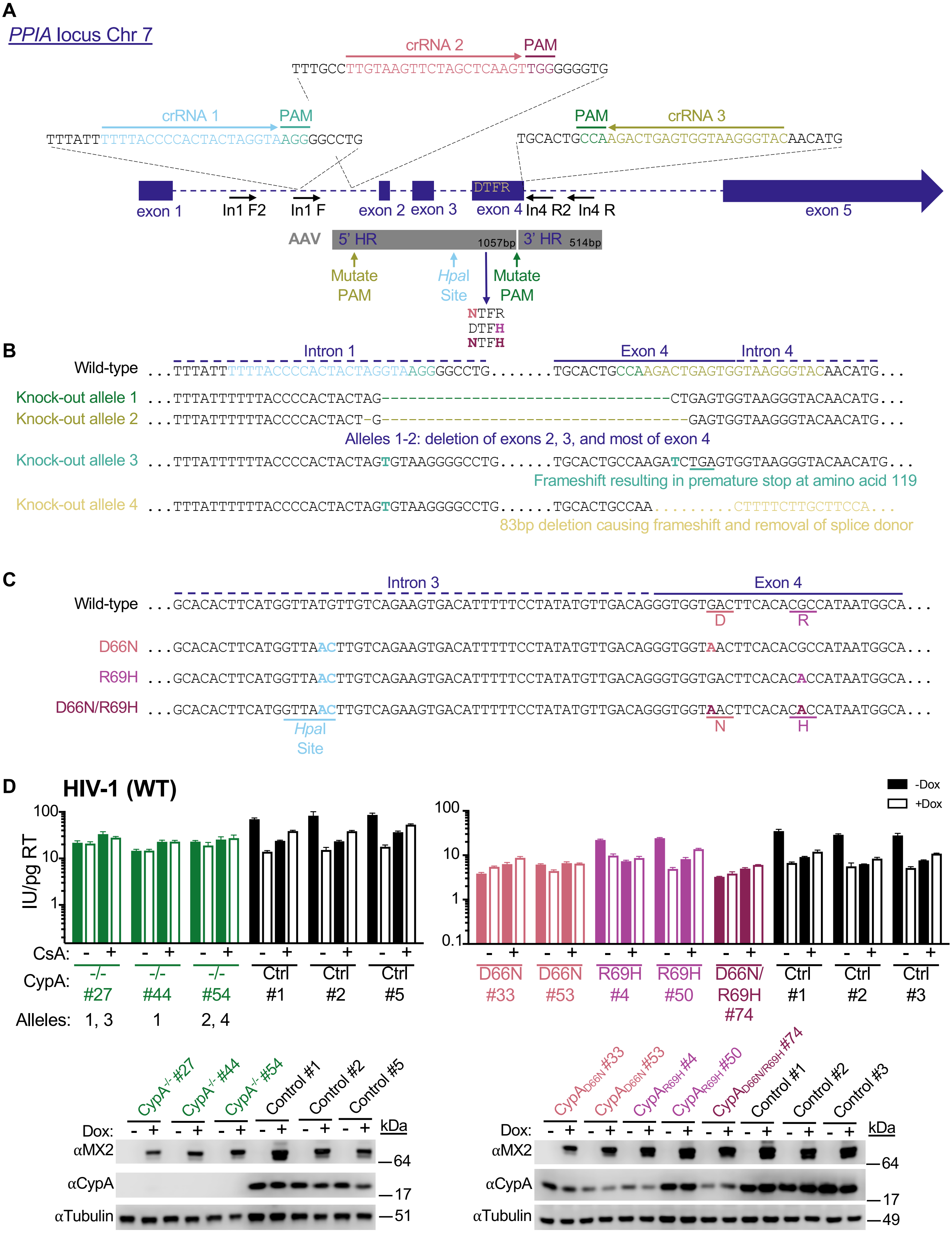


**S2 Fig. Generation of CypA knockout and mutant cell lines**

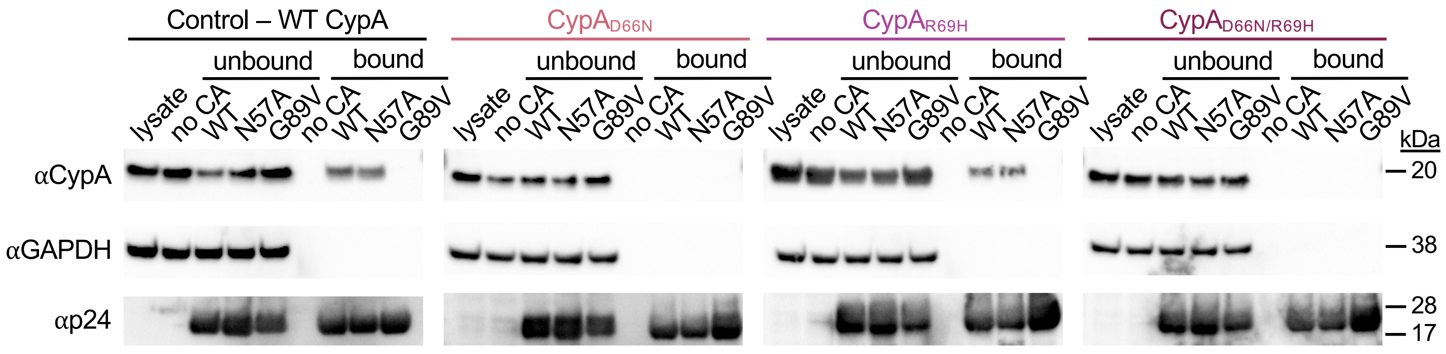


**S3 Fig. Validation of CA binding specificity of CypA mutants**

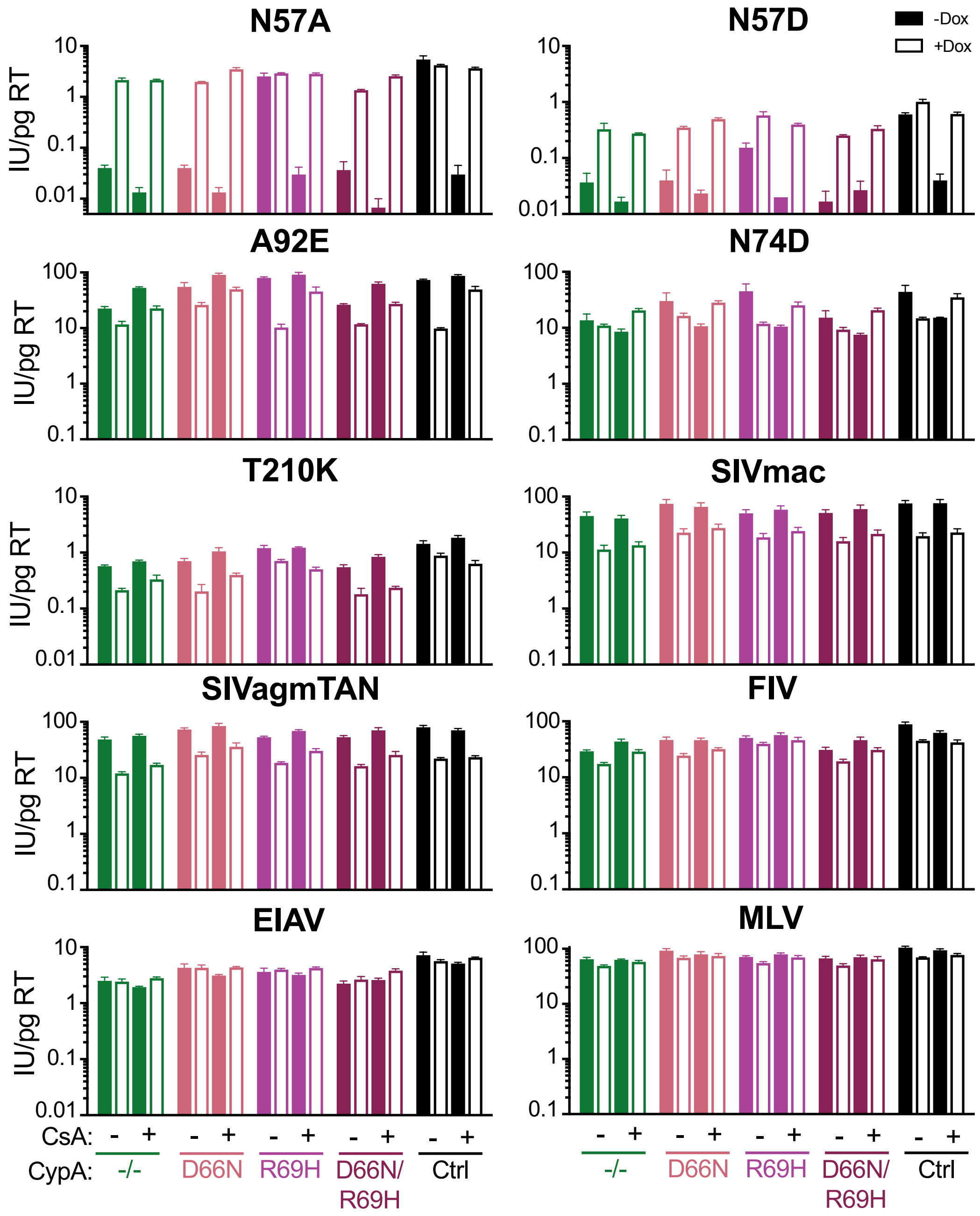


**S4 Fig. The effects of CsA on retroviral sensitivity to MX2 are the direct result of blocking CA-CypA interactions**

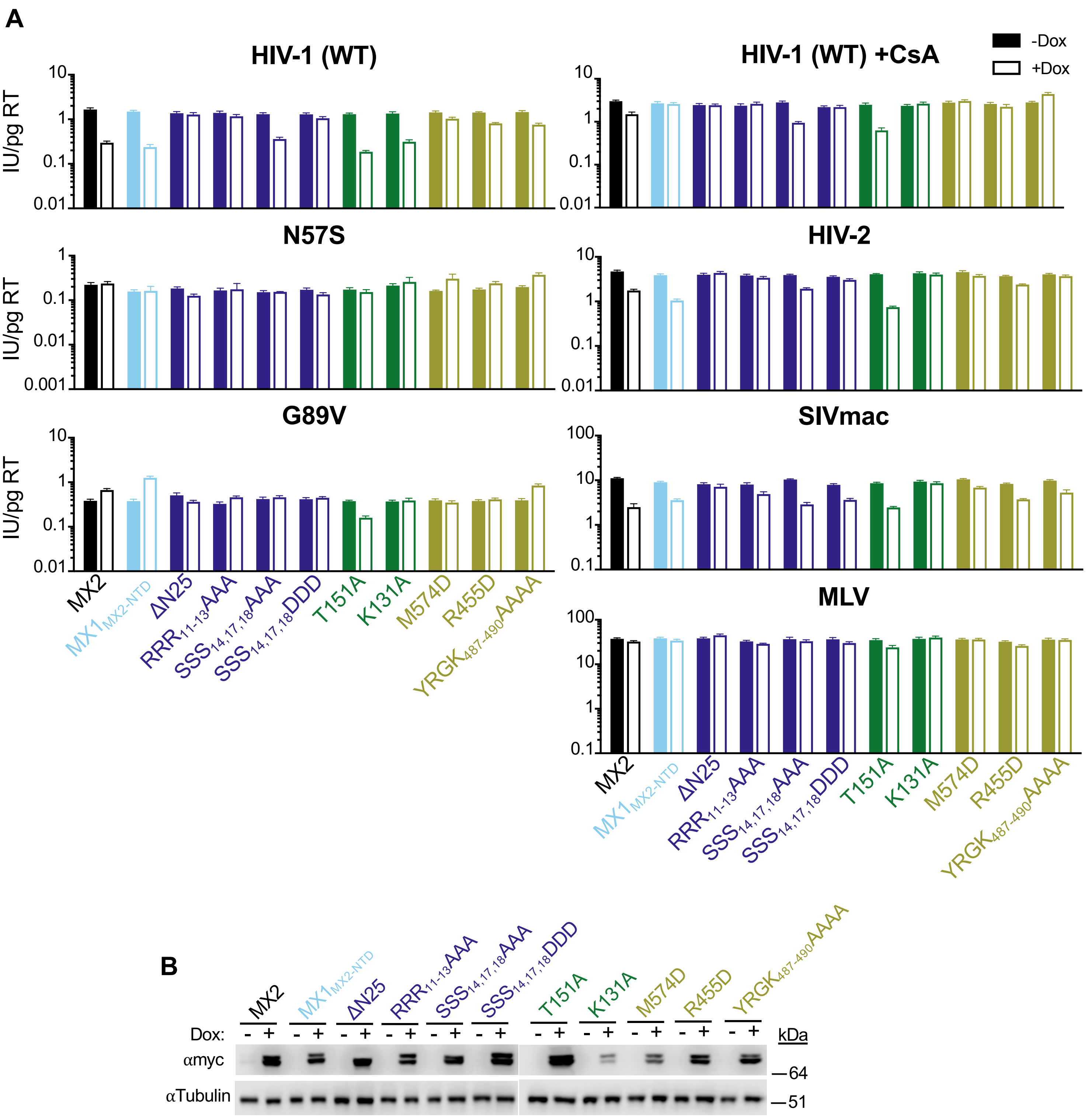


**S5 Fig.– Determinants for MX2 activity in the absence of CA-CypA interactions**

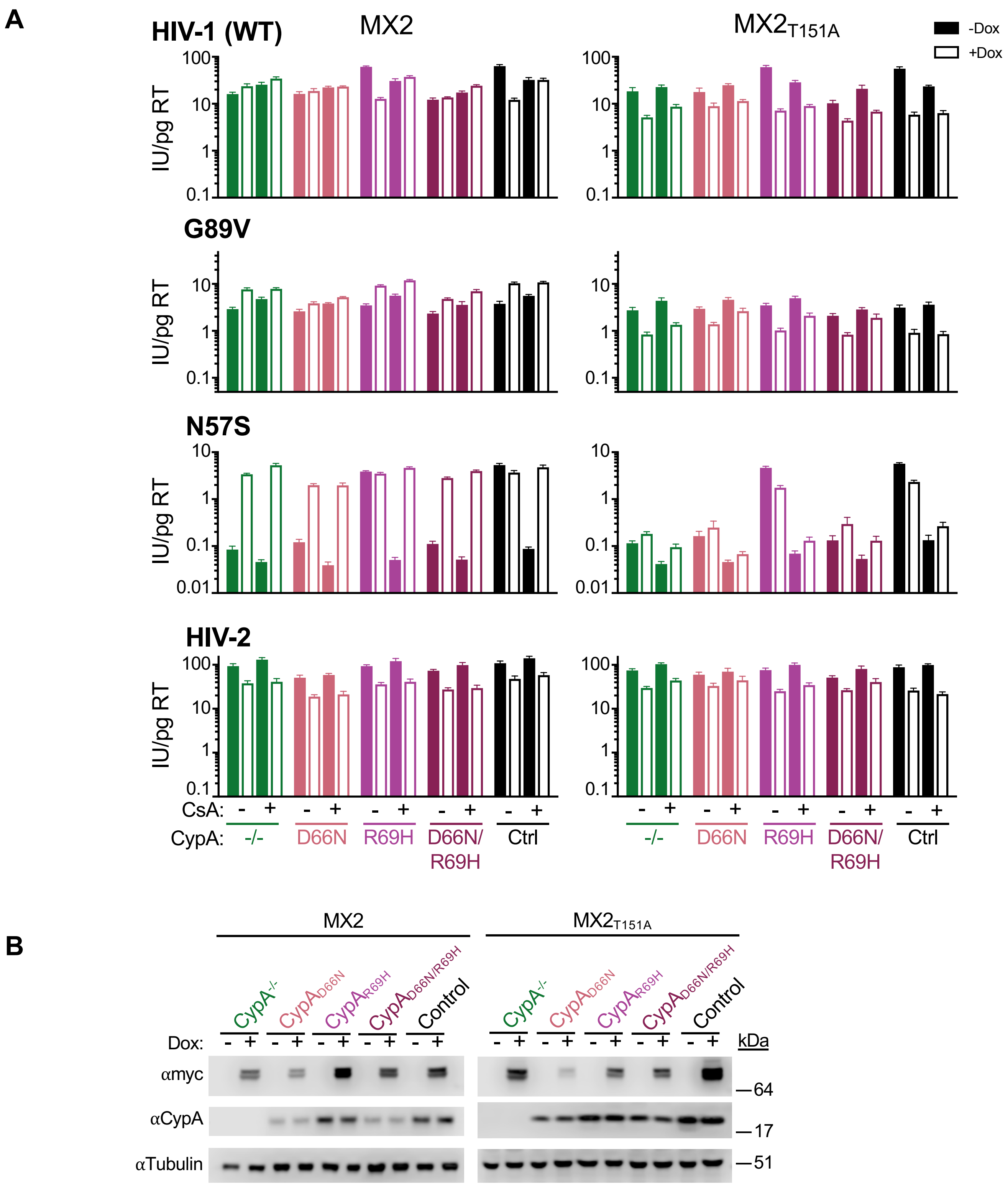


**S6 Fig. – Restriction of HIV-1 infection by GTPase-deficient MX2 in CypA mutant cells**

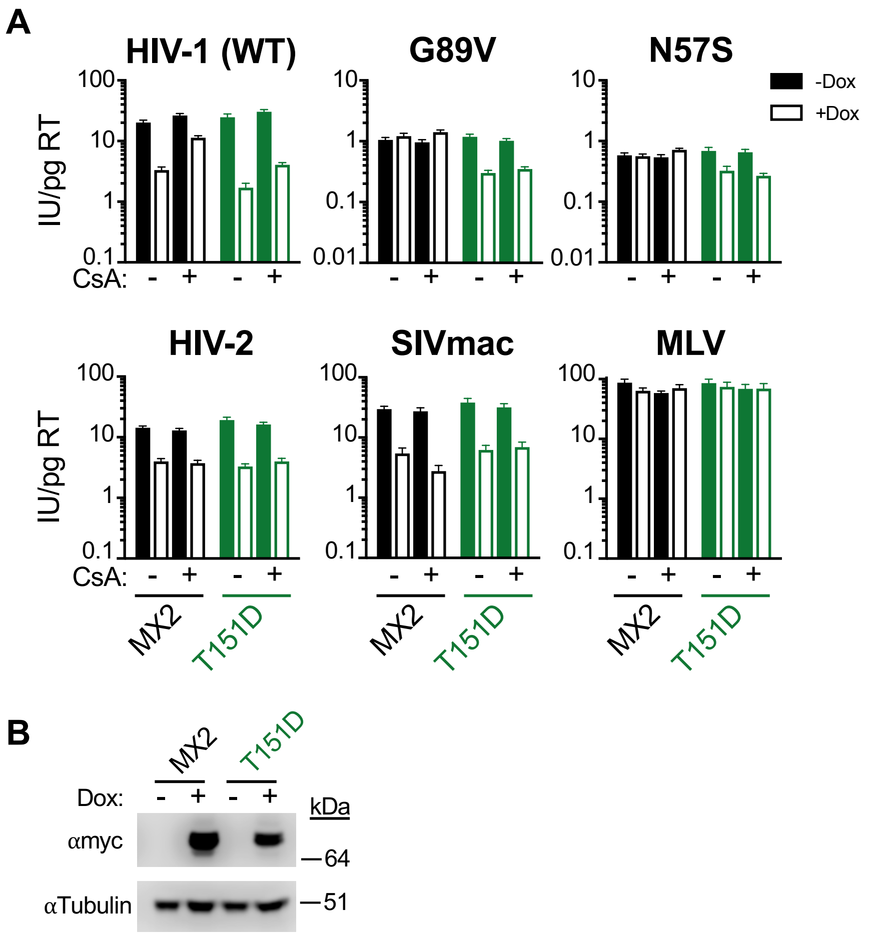


**S7 Fig – Phosphorylation at residue T151 does not determine CA-CypA-dependent MX2 activity**

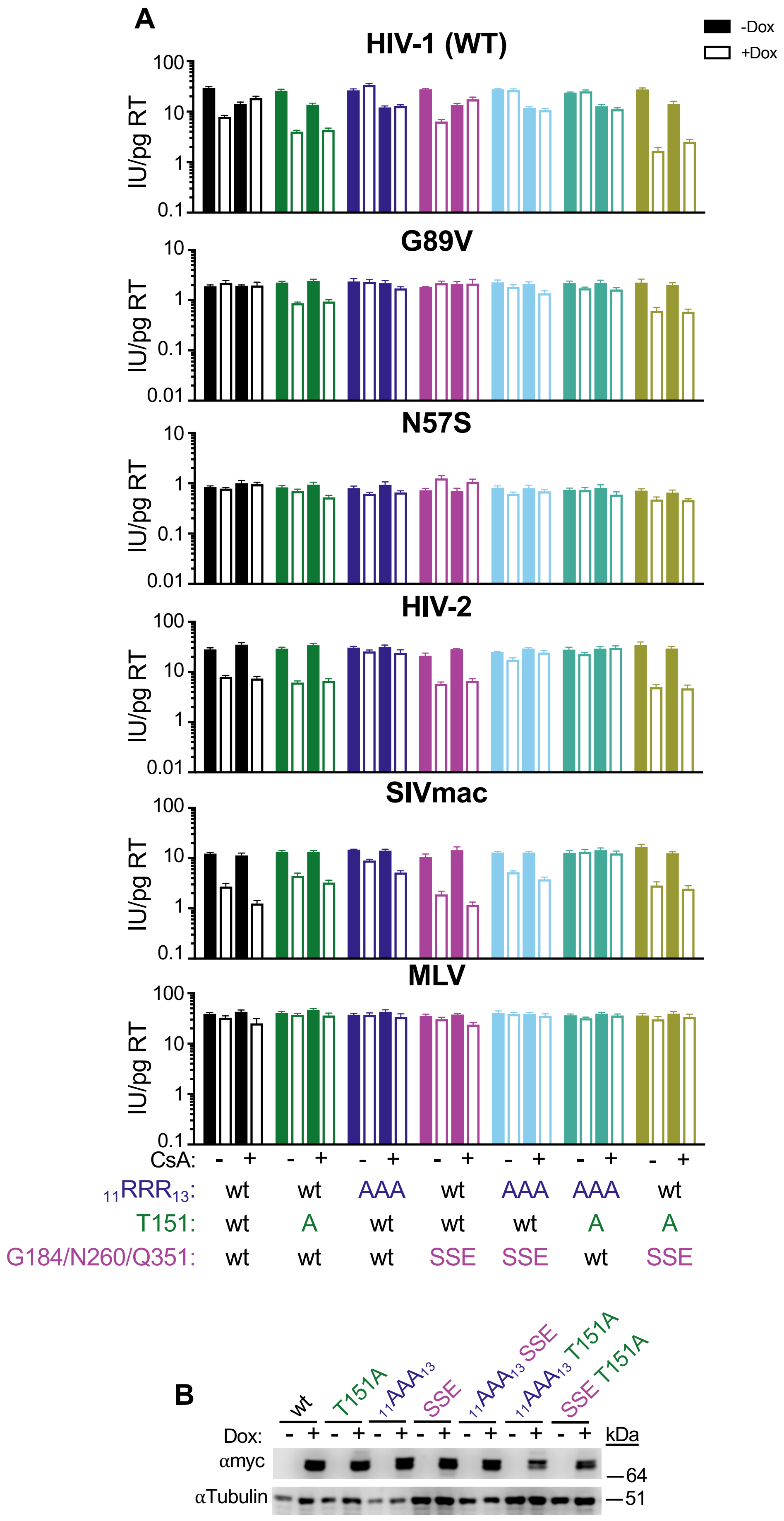


**S8 Fig – Restriction by GTPase-deficient MX2 in the absence of CA-CypA binding is not mediated by known CA-GTPase domain interactions**

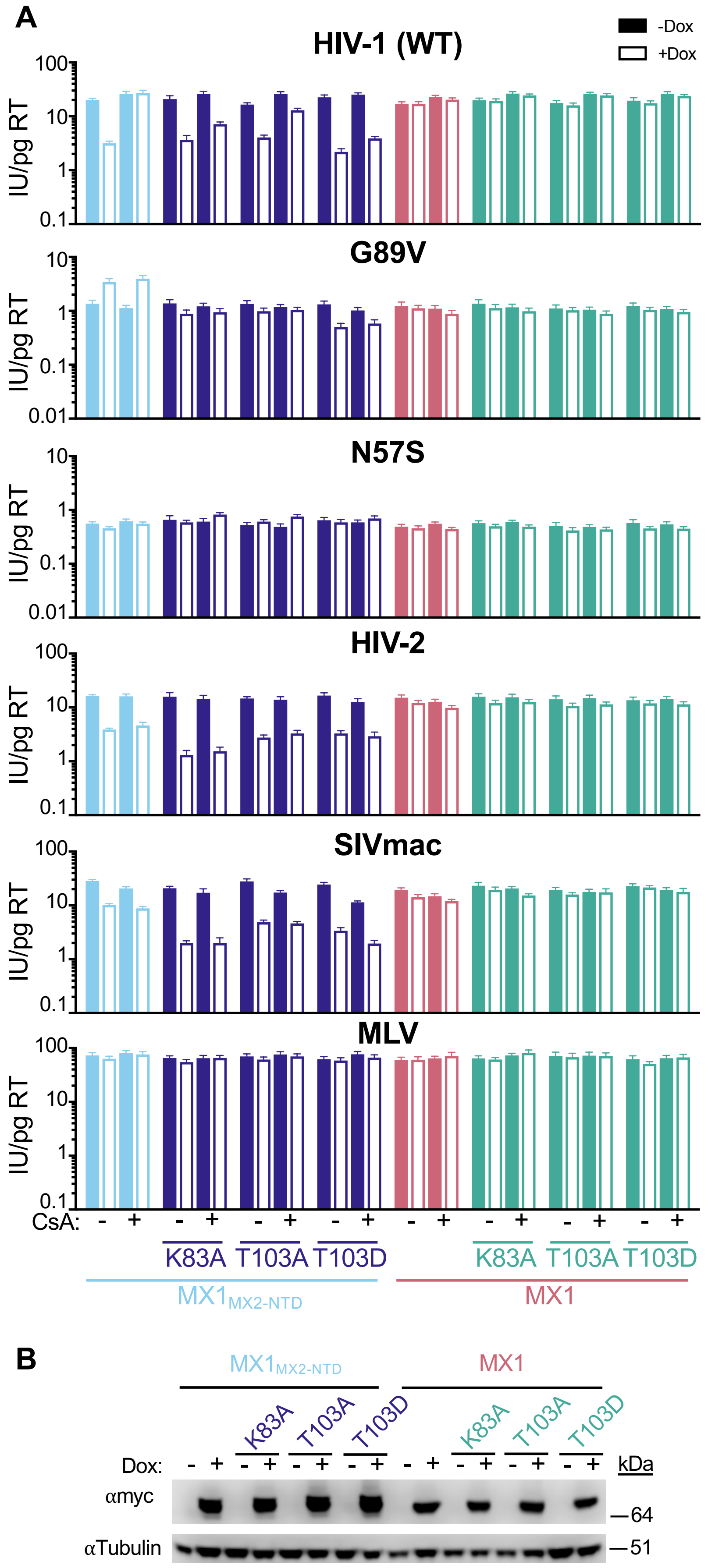


**S9 Fig – Antiviral activity of GTPase-deficient chimeric $MX1_{MX2\text{-}NTD}$ proteins**

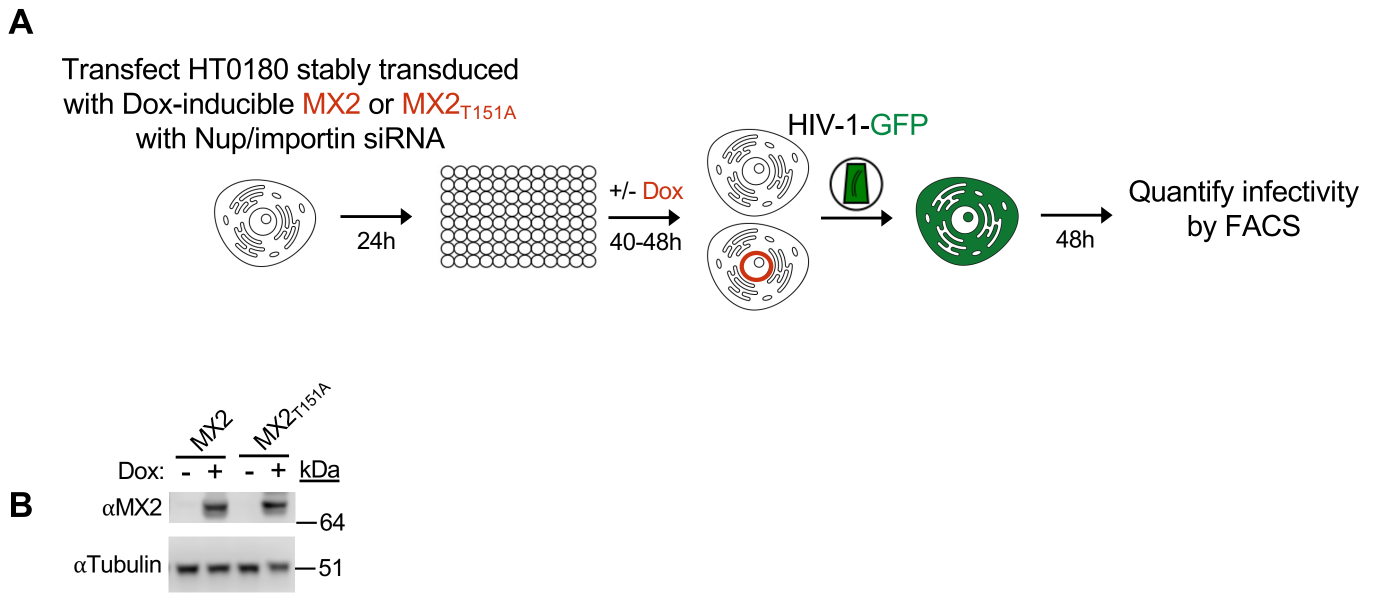


**S10 Fig – Experimental design for Nup/NTR knockdown**

### S1 Table. PCR primers

| | |
|---|---|
| **MX2 mutants** | |
| Mx2-SfiI-F | CTCTGGCCGAGAGGGCCATGTCTAAGGCCCACAAGCCTTG |
| Mx2-myc-SfiI-R | CTCTGGCCAGAGAGGCCTCACAGATCCTCTTCAGAGATGAGTTTCTGCTCGTGGATCTCTTTGCTGG AGAA |
| Mx2-SfiI-R | CTCTGGCCAGAGAGGCCTCAGTGGATCTCTTTGCTGGAG |
| Mx2-M574D-F | TGATCCAACTTCAGTTCAGAGACGAGCAGATGGTTTTTTGTC |
| Mx2-M574-R | GACAAAAAACCATCTGCTCGTCTCTGAACTGAAGTTGGATCA |
| Mx2-Y651D-F | CCAGATCCCATTTATAATTCAGGATTTTATGCTCCGAGAGAATGGTGACTCC |
| Mx2-Y651D-R | GGAGTCACCATTCTCTCGGAGCATAAAATCCTGAATTATAAATGGGATCTGG |
| Mx2-YRGK-AAAA F | GTTGAAAAATATGAAAAGCAGGCTGCAGCCGCGGAGCTTCTGGGATTTGTCAAC |
| Mx2-YRGK-AAAA R | GTTGACAAATCCCAGAAGCTCCGCGGCTGCAGCCTGCTTTTCATATTTTTCAAC |
| Mx2-RRR-11-13-AAA-SfiI-F | ctctggccgagagggccATGTCTAAGGCCCACAAGCCTTGGCCCTACGCGGCGGCAAGT |
| Mx1-Mx2N91-F | CGAGAACAACCTGTACAGCCAGTATGAGGAGAAGGTGCGC |
| Mx1-Mx2N91-R | GCGCACCTTCTCCTCATACTGGCTGTACAGGTTGTTCTCG |
| Mx1-nostop-SfiI-R | ctctggccagagaggccACCGGGGAACTGGGCAAGCCG |
| Mx1-SfiI-F | ctctggccgagagggccATGGTTGTTTCCG |
| Mx2-R455D-F | CCCGTTTATACAACAAAATCGACGAGGATTTTAAAAACTGGG |
| Mx2-R455D-R | CCCAGTTTTTAAAATCCTCGTCGATTTTGTTGTATAAACGGG |
| Mx2-delN25-SfiI-F | ctctggccgagagggccATGAATTCCTTCC |
| Mx1-myc-Sfi1-R | ctctggccagagaggccTCACAGATCCTCTTCAGAGATGAGTTTCTGCTCACCGGGGAACTGGGCAAGCCG |
| Mx2-SfiI-RRR-AAA-F2 | CTCTGGCCGAGAGGGCCATGTCTAAGGCCCACAAGCCTTGGCCCTACGCGGCGGCAAGTCAATTTT CTTCTCG |
| Mx2-SfiI-SSS-AAA-F | ctctggccgagagggccATGTCTAAGGCCCACAAGCCTTGGCCCTACCGGAGGAGAGCTCAATTTGCTGCT CGAAAATACCTG |
| Mx2-SfiI-SSS-DDD-F | ctctggccgagagggccATGTCTAAGGCCCACAAGCCTTGGCCCTACCGGAGGAGAGATCAATTTGATGATC GAAAATACCTG |
| Mx2-T151D-F | GCGGAATCGTAGACAGGTGTCCGC |
| Mx2-T151D-R | GCGGACACCTGTCTACGATTCCGC |
| Mx1-K83A-F | CCAGAGCTCGGGCGCGAGCTCCGTG |
| Mx1-K83A-R | CACGGAGCTCGCGCCCGAGCTCTGG |
| Mx1-T103A-F | GCGGGATCGTGGCCAGATGCCCG |
| Mx1-T103A-R | CGGGCATCTGGCCACGATCCCGC |
| Mx1-T103D-F | GCGGGATCGTGGACAGATGCCCG |
| Mx1-T103D-R | CGGGCATCTGTCCACGATCCCGC |
| **TRIM5-fusions** | |
| CypA R69H F | CTTCACACACCATAATGGCACTGGT |
| CypA R69H R | ACCAGTGCCATTATGGTGTGTGAAG |
| CypA NotI F | ataagaatgcggccgccATGGTCAACCCCACCGTG |
| CypA SalI R | ataagaatgtcgactcaTTCGAGTTGTCCACAGTCAGC |
| CypA D66N F | GGTGGTAACTTCACACGCCATAATG |
| CypA D66N R | CATTATGGCGTGTGAAGTTACCACC |
| CypA (NH) F | GGTGGTAACTTCACACACCATAATG |
| CypA (NH) R | CATTATGGTGTGTGAAGTTACCACC |
| **AAV** | |
| AAV NheI F | TTCCTGCGGCCGCAGCATAGCTAGC |
| AAV XhoI R | CGCTCGGTCCGCACAATTCCTCGAG |
| **HIV-1 CA mutants** | |
| CapNM F | GTA AGA AAA AGG CAC AGC AAG CGG CCG CTG |
| CapNM R | CTT GGC TCA TTG CTT CAG CCA AAA CGC GTG |
| N57A F | AACACCATGCTAGCCACAGTGGGG |
| N57A R | CCCCACTGTGGCTAGCATGGTGTT |
| N57D F | AACACCATGCTAGACACAGTGGGG |
| N57D R | CCCCACTGTGTCTAGCATGGTGTT |
| **CypA-mutant cell verification** | |
| CypA Intron1 F2 (In1 F2) | TCTAAACTTGGCGCGTGTCT |
| CypA Intron 4 R (In4 R) | TTCAACCACCCAGCTAAGGG |
| CypA Intron 1 F (In1 F) | AAGAGAAGTGCACACGGATACT |
| CypA Intron 4 R2 (In4 R2) | GTCAGGTGGTTAGTGTGCCA |
